# BinSanity: Unsupervised Clustering of Environmental Microbial Assemblies Using Coverage and Affinity Propagation

**DOI:** 10.1101/069567

**Authors:** Elaina Graham, John Heidelberg, Benjamin Tully

## Abstract

Metagenomics has become an integral part of defining microbial diversity in various environments. Many ecosystems have characteristically low biomass and few cultured representatives. Linking potential metabolisms to phylogeny in environmental microorganisms is important for interpreting microbial community functions and the impacts these communities have on geochemical cycles. However, with metagenomic studies there is the computational hurdle of ‘binning’ contigs into phylogenetically related units or putative genomes. Binning methods have been implemented with varying approaches such as k-means clustering, Gaussian mixture models, hierarchical clustering, neural networks, and two-way clustering; however, many of these suffer from biases against low coverage/abundance organisms and closely related taxa/strains. We are introducing a new binning method, BinSanity, that utilizes the clustering algorithm affinity propagation (AP), to cluster assemblies using coverage alone, removing potential composition based biases in clustering contigs, but requires a minimum of two samples. To increase fidelity, a refinement script was developed that uses composition data (tetranucleotide frequency and %G+C content) to refine bins containing multiple source organisms. This separation of composition and coverage based signatures reduces clustering bias for closely related taxa. BinSanity was developed and tested on artificial metagenomes varying in size and complexity. Results indicate that this implementation of AP lead to a higher precision, recall, and Adjusted Rand Index over five commonly implemented methods. When tested on a previously published infant gut metagenome, BinSanity generated high completion and low redundancy bins corresponding with the published metagenome-assembled genomes.

## Introduction

Studies in microbial ecology commonly experience a bottleneck effect due to difficulties in microbial isolation and cultivation. This bottleneck has long since been acknowledged through comparisons of direct microscopy and culturing^1^. Despite the culture bottleneck, research into different environments reveals a higher diversity of organisms than had been previously estimated. Much of this research occurs through sequencing the collective genomes (metagenome) of microorganisms in the environment^2^. Using metagenomics allows for the linkage of mechanistic pathways, metabolism, and taxonomy to infer environmental context without cultivation. Due to the decrease in sequencing cost, assembly and binning have become two major bottlenecks for conducting metagenomics research due to large computational requirements and methods with limited fidelity. Recent advances have decreased the limitations of metagenomic assemblers, but binning remains a second major computational bottleneck in metagenomics. Typically, one of a few issues are encountered in current binning protocols, including: decreasing accuracy for contigs below a size threshold, necessity of human intervention in distinguishing clusters, struggling to differentiate related microorganisms with similar k-mer frequencies, or excluding low coverage and low abundance organisms.^3-5^

Popular unsupervised binning methods commonly use compositional parameters, such as tetranucleotide frequency^6,7^ as the major delimiting parameter for creating putative groups of related sequences (bins). Due to the taxon specific nature of codon usage^8,9^ GC content^8,10^, and short oligonucleotides (k-mers)^11,12^ these useful fingerprints have been used to characterize and cluster contigs.

However, the utilization of composition alone can lead to biases during binning due to closely related species having similar fingerprints, as well as, recently acquired genes from horizontal transfer creating aberrant connections^13^. These issues could lead to incomplete and overly chimeric organisms. Several methods and protocols have had increased success by incorporating coverage information as an additional variable for determining bins. Development of new binning protocols are essential to better characterize complex environmental communities and explore microbial diversity at a level that cultivation based studies at present cannot achieve.

BinSanity utilizes the clustering algorithm Affinity Propagation (AP) and accepts contig coverage values as the primary delimiting component. We propose that BinSanity is a more accurate alternative to previous clustering implementations. Further, while other clustering algorithms can effectively group related DNA fragments using composition and optional incorporation of coverage data, common methods, like hierarchical clustering and k-means require human input of information criteria that dictate the ultimate number of clusters (e.g Bayesian information criterion). Accurately representing the community diversity in this number is increasingly difficult as community complexity increases. AP, in contrast, requires no input on determining cluster centers; instead every point is iteratively considered as a potential cluster center. Within BinSanity, each contig is evaluated as a possible exemplar based on the coverage. The exemplar is the contig that best represents the contigs clustering with it and can also be referred to as the cluster center. AP is described elsewhere^14-16^, but in brief, AP takes as input a collection of values where the similarity ***s*(*i*,*k*)** indicates how well the data point with index ***k*** is suited to be the exemplar for data point ***i***. The messages sent between points make up either the responsibility ***r*(*i*,*k*)** or the availability ***a*(*i*,*k*)**^16,17^. The responsibility is the accumulated evidence that sample ***i*** should be the exemplar for sample ***i*** (Formula 1)^15^. The availability^15^ is the accumulated evidence that sample ***i*** should choose sample ***k*** to be its exemplar, dually considering the evidence of values for other samples that ***k*** should be an exemplar (Formula 2). Two limitations of AP are that it is hard to pinpoint the optimal preference (p) and damping factor. The preference is a measure of whether data point ***i*** should be chosen as an exemplar. High values of a preference will lead to more exemplars (splitting) and low preferences will lead to a smaller number of exemplars (lumping). When setting a global value for AP, the minimum similarity is typically used as an initial choice for the preference^16^. The damping factor is a number that helps to account for exemplars in periodic variance during the iterative process as well as improves convergence during oscillations^18,19^. In addition, AP faces the challenge of time and memory complexity in the order of ***o*(*N^2^T*)** where N is the number of samples and T is this number of iterations until convergence^14-16,18^. This order does not scale for production of a dense similarity matrix.

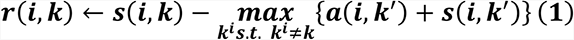

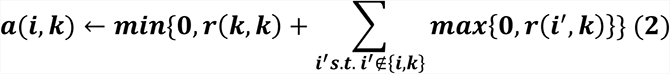

Data shows that AP effectively clusters a variety of data types and is often more precise than similar clustering methods^14-17,19-23^. Another implementation of AP for clustering contigs was developed by Lin and Liao ^24^ (myCC), where a two stage approach utilized single copy marker genes, tetranucleotide frequencies, and the optional input of coverage. BinSanity, in contrast, bypasses composition based biases on clustering by creating an initial set of clusters using coverage and optionally refining with composition when coverages of multiple organisms converge. The ability to bin less contaminated and more diverse taxa from metagenomics data will enhance our ability to uncover what microbial groups are present and the corresponding putative biogeochemical processes.

We benchmarked BinSanity by comparing it to the current generation of unsupervised binning software. We constructed several artificial microbial communities and created *in silico* metagenomic samples based on these sequences. The communities were composed of sequences that can be problematic for composition based binning algorithms, specifically metagenomes consisting of closely related and low abundance organisms. Additionally, a dataset associated with an infant gut microbiome time-series was used to establish how clusters generated via BinSanity compared to a highly curated set of genomic bins originally constructed using a manual ESOM^13^ based.

## Methodology

### Artificial Metagenomes

In total 60 reference genomes (including some closed genomes, some metagenome-assembled genomes [MAGs], and some draft genomes; Supplemental Table S1) consisting of a variety of organisms with ecological and environmental significance were accessed from the Joint Genome Institute (JGI) Integrated Microbial Genome (IMG) Portal^25^ and NCBI^26^ and used to create *in silico* microbial communities. Reference genomes were screened via CheckM^27^ to provide a baseline for estimates of completion and redundancy. For each community, *in silico* metagenomes were generated using the reads-for-assembly script (https://github.com/meren/reads-for-assembly), which generates “sequence reads” from the source DNA that mimics random variations around an assigned coverage value and generates reads with simulated next-generation sequencing lengths and error rates. Because the script simulates variations around a mean-coverage value, genomes with assemblies greater than 20kbp (or closed genomes) were randomly split in to fragments between 3kbp and 15kbp in length using a Python script (split_file.py). For each metagenome, organisms were assigned to be either low (randomly assigned a coverage value <10X) or high abundance (randomly assigned a coverage value between 10X-200X) by an in-house script (make_config_ini.py).

Three artificial communities were constructed to test BinSanity. The first artificial community selected 25 organisms of the original 60 reference genomes, including four strains of *Escherichia coli* (further referenced as, strain-mixture) and the organisms were randomly assigned as low (n = 13) or high abundance (n = 12); low and high abundance organisms were randomly reassigned for each metagenome. Organisms could alternate between low and high abundance categories between samples. Two additional communities with 50 organisms from distinct species were curated from the 65 reference genomes. One of the communities had half of the organisms (n = 25) randomly assigned to either be in either low or high abundance for each metagenomic sample (diverse-mixture-1). In the other community, all of organisms were assigned to be low abundance (diverse-mixture-2).

After the reads for each *in silico* metagenome were generated, the reads were aligned back to the reference genomes using Bowtie2^28^ (v2.2.5; default parameters). The output SAM file was then converted to a BAM file using SAMtools^29^ (v1.2 parameters: samtools view -bS file | samtools sort - file). This BAM file was used to calculate the coverage for each contig (reads/bp) via an in-house script (contig-coverage-bam.py) that implements BEDtools^30^. The determined coverage values were log transformed and results from multiple metagenomes were combined in to a single matrix using an in-house script (cov-combined.py).

Genomes were reconstructed for each of the three artificial metagenomes based on the binning results utilizing coverage values for 20, 15, 10, 5, 4, 3, and 2 *in silico* metagenomes. BinSanity was executed on the log transformed coverage matrix using the script BinSanity.py (-m 4000 -v 400 -d 0.95 -x 1000 -p [variable]; for low abundance samples, other modifications to coverage value can be applied, e.g. transforming coverage values by a set value to increase resolution, etc.). For the strain-mixture, a preference of -3000 was used (-p -3000). For diverse-mixture-1, 2-5 *in silico* metagenomes used a preference of -5 (-p -5); 15 and 20 metagenomes used a preference of -1000 (-p -1000). For the diverse-mixture-2, the preference was set between -10 and -20 depending on the number of *in silico* metagenomes (20 and 15 *in silico* metagenomes -p = -20; 10 *in silico* metagenomes -p = -15; 2-5 *in silico* metagenomes -p = -10). For the infant gut metagenome preference was set to -10 (-p -10). Changes in the preference value are discussed below. BinSanity was compared against CONCOCT^3^ (v.0.4.1; default parameters), GroopM^5^ (v0.3.5; default parameters), MetaBat^31^ (v0.26.3; default parameters), and MaxBin32 (v2.1.1; default parameters). All of the compared methods use both composition and coverage to determine bins, though coverage is optional for all, except GroopM. BinSanity could be used with or without composition information. For the purposes of our analysis, we used a composition based refinement function (BinSanity-refine.py; -m 4000 -v 400 -x 1000 -d 0.95 -p -500) to refine bins with high redundancy. This script uses percent G+C (G+C%) content, coverage, and tetranucleotide frequencies.

Results were evaluated by calculating precision, recall, and V-measure ( *e.g.* harmonic mean) as defined by Rosenberg and Hirschberg ^33^ using sklearn.metrics.homogeneity_completeness_v_measure^34^ (bin_evaluation.py). Precision defines whether each cluster contains only members of a single class (an output of 1 representing all bins contain only contigs from a single source). Recall considers whether each member of a class is assigned to the same bin (an output of 1 representing that only contigs from one source organism are contained in a single bin). The V measure is the harmonic mean of the precision and recall allowing evaluation of accuracy. An additional measure, the Adjusted Rand Index (ARI)^35^ was also calculated via sklearn.metrics.adjusted_rand_score^34^ (bin_evaluation.py). The ARI considers similarity between predicted and true cluster labels. This similarity is then adjusted for chance using a probability heuristic. In addition, analysis of the clustering results via CheckM^27^ for completion, redundancy, and strain heterogeneity was also conducted. The general workflow for affinity-propagation is shown in Figure 1.

**Figure 1.**
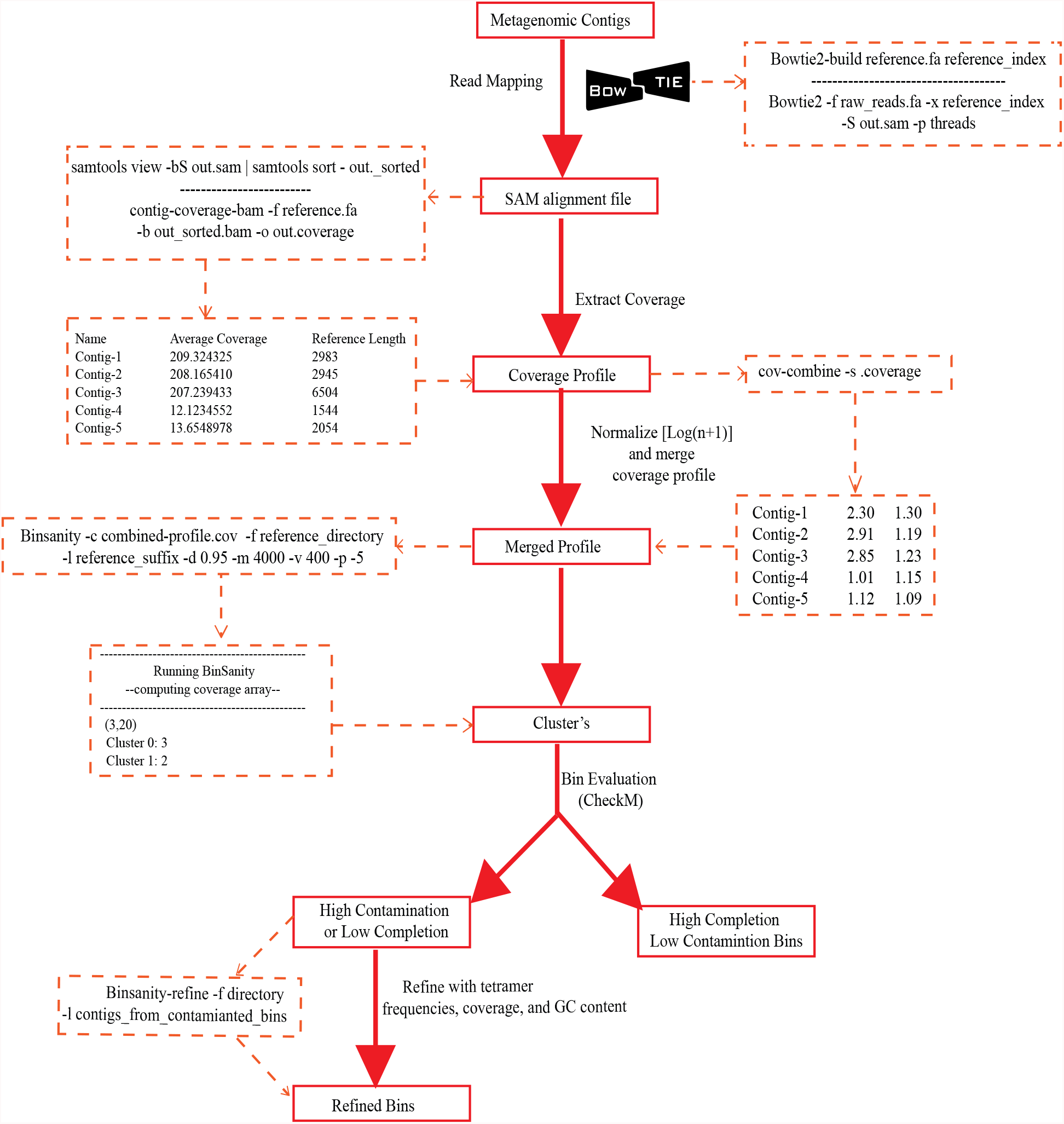
Workflow for Binsanity indicating all scripts used.

### Infant Gut Metagenome

BinSanity was assessed using real data, using samples from a time series study of an infant gut microbiome, previously described by Sharon, et al. ^36^. Samples were run though BinSanity.py using the following parameters: -p -10 -m 4000 -v 400 -d 0.95 -x 100. This same dataset was accessed by Eren, et al. ^37^ and was binned using a human guided strategy via the Anvi’o platform^37^. In an effort to measure the effect of the binning algorithms (and to avoid influencing the results due to the use of different assemblers) the contigs produced by Eren, et al. ^37^ (http://anvio.org/data/) were used as the input for BinSanity (Eren-contigs). Raw reads were accessed from the NCBI SRA database (SRA052203) and aligned to the Eren-contigs and the coverage matrix was determined as described above. The Eren-contigs were also binned using CONCOCT, GroopM, MaxBin, and MetaBat. All genome bins were evaluated via CheckM^27^ and compared to genomes generated by Sharon, et al. ^36^ (http://ggkbase.berkeley.edu/carrol/). To maintain consistency, the curated bins from Sharon, et al. ^36^ were processed using CheckM, so that all genome bin metrics were consistent.

## Results and Discussion

### Species Level: Diverse-Mixture-1

In processing diverse-mixture-1, BinSanity had near perfect results generating 50 bins (Figure 2) with an ARI and V-measure of 0.98 using 20 *in silico* metagenomes (Figure 3). When the number of *in silico* metagenomes was decreased to five, BinSanity still had the highest ARI and V-measure. With five *in silico* metagenomes, BinSanity produced a total of 62 bins, 43 of which were >90% complete (as determined by CheckM). Of the remaining seven organisms, five were >70% complete. The remaining contigs were contained in small bins, only one of which contained contigs from multiple sources. Two genomes were each split into two bins (for a total of four); each with no contamination. BinSanity produced the highest V-measure score of the binning methods, indicating it most closely reconstructed the reference organisms and had minimal rates of incorrectly assigned contigs. Additionally, the completion and redundancy for the 43 bins >90% complete were closer to the expected values from the stock organisms than any of the other four methods (Supplemental Table S2).

**Figure 2.**
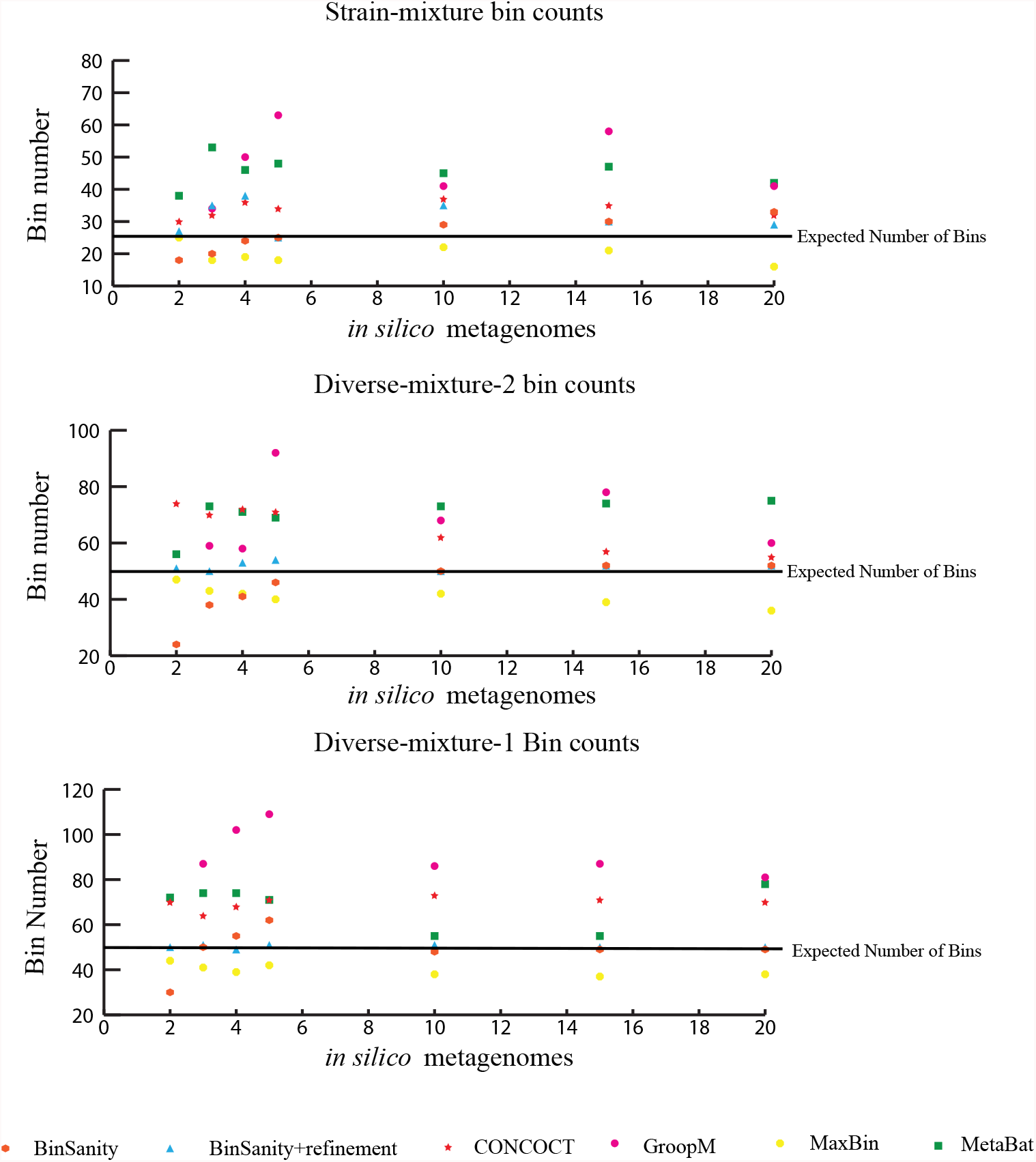
Each graphs shows the raw number of bins output from each method at each in silico metagenome number tested (e.g 2,3,4,5,10,15,20). The black line represents the expected number of bins. In the strain mixture raw bin counts oscilated between being above and below the expected relative to the number of in silico metagenomes used. Adding composition information via the refinement script minimally impacted bin number betwee 10 and 20 in silico metagenomes for Diverse-mixture-2 and Diverse-mixture-1. In the Strain-mixture incorporation of composition via the refinement script and Binsanity lead to increased numbers of bins.

**Figure 3.**
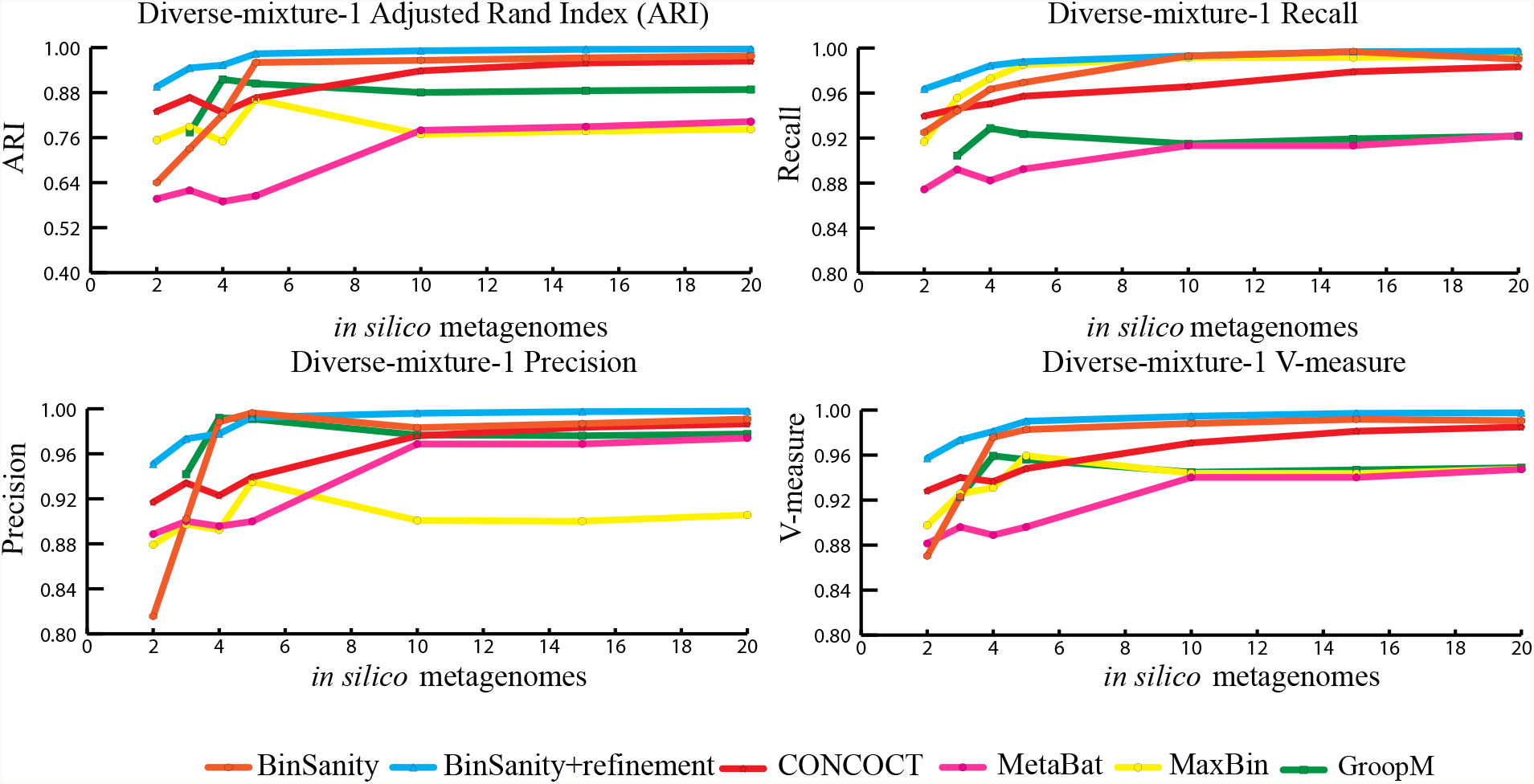
stastistical calculations (bin_evaluation.py) showing the Adjusted rand index (ARI), Precision, Recall, and V-measure for diverse-mixture-1 which included 25 rarebiosphere and 25 abundant biosphere organisms. Above 4 in silico metagenome BinSanity yielded the highest ARI, Precision, and V-measure. Below 4 in silico metagenomes CONCOCT and GroopM had the highest ARI, Precision, and V-measure. When BinSanity was used with refinement the script, it was able to maintain the highest ARI, precision, and V-measure below 4 in silico metagenomes.

In comparison to BinSanity, GroopM and MetaBat had high precision but low recall (indicating too many bins were produced), whereas CONCOCT and MaxBin had high recall but a low precision (indicating too few bins were produced). With an expected output of 50 bins at five *in silico* metagenomes, CONCOCT, GroopM, MaxBin, and Metabat produced 71, 109, 47, and 72 bins, respectively. While BinSanity produced 43 bins at >90% complete, MetaBat and GroopM produced 33 and 41, respectively. Of the 47 bins produced by MaxBin, eight were highly chimeric (Figure 4). CONCOCT, overall, had a high accuracy, but had difficulty delimiting closely related species such as *Roseobacter denitrificans* and *R. litoralis*. This difficulty in separating closely related species could be related to the use of a single step clustering protocol where composition and coverage are used as equally weighted inputs. Closely related organisms often have similar composition based signals (G+C%, tetranucleotide frequency, etc.). Coverage, in contrast, is reliant on the underlying population of the organisms in question, such that contigs from the same organisms should have closely related coverage values. One caveat to coverage based methods is that reads need to be assigned to contigs with minimal bias for conserved regions and nonspecific alignment. Strict alignment parameters (such as using the -- very-sensitive flag in Bowtie2) can be used to prevent false contig assignments and increase fidelity of all of the binning methods. And more input coverage information, especially variable coverage data, benefits all of the methods, as is evident when analyzing results generated using <5 *in silico* metagenomic samples; all methods decline in accuracy.

**Figure 4.**
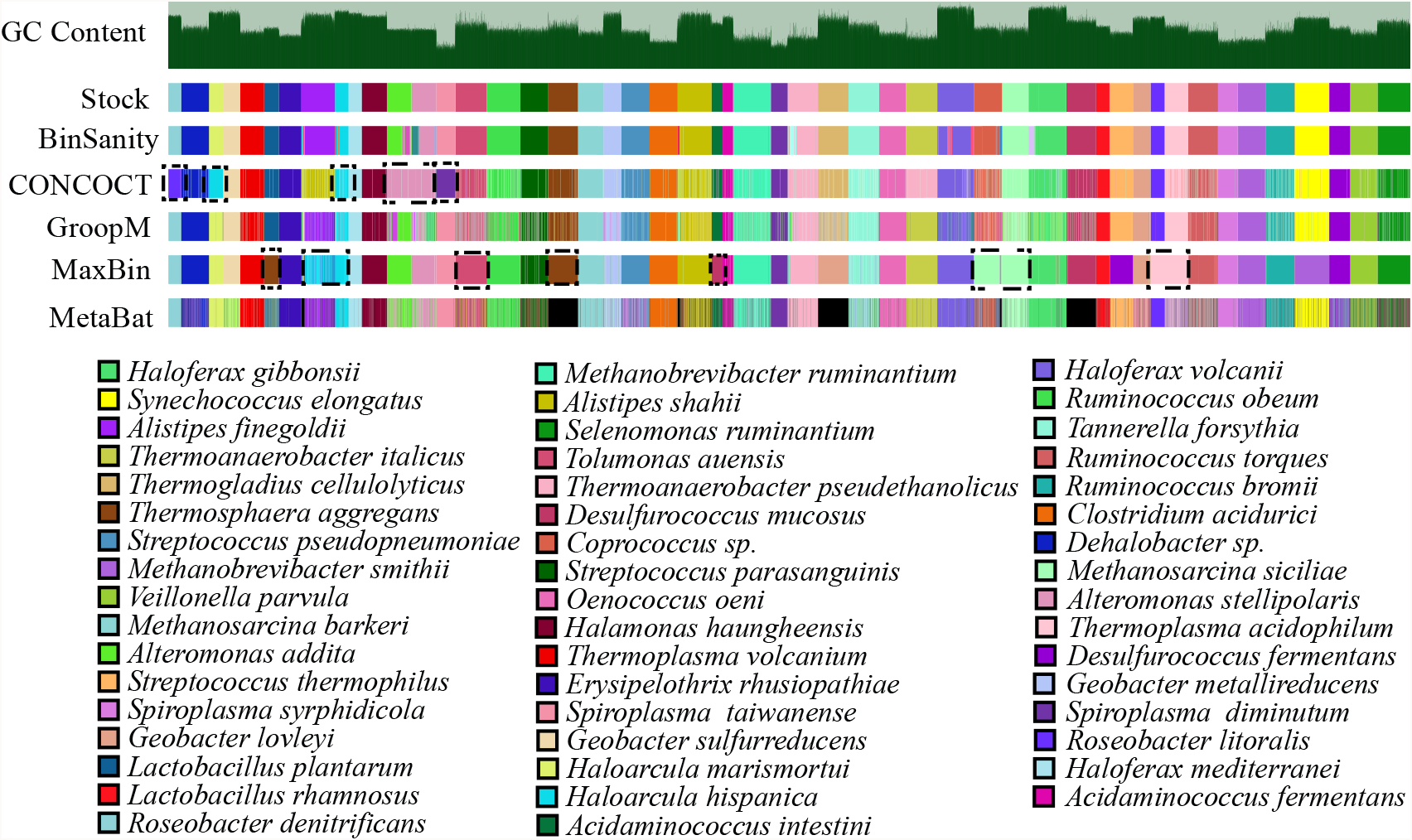
Image generated via Anvi’o. Exhibits clustering of diverse-mixture-1 at five in silico metagenomes. Contigs were ordered based on coverage and organism designation from the stock mixture shown first. Black was used to emphasize contigs that remained unbinned in the final results. Black dashed boxes outline organisms that were clumped together by the clustering methodi n question. AP had no issues at 5 in silico metagenomes with over clumping whereas CONCOCT, GroopM, and MaxBin did. MetaBat on the other hand left the most contgis unclustered.

The primary method for generating bins within BinSanity is clustering using coverage values. When the number of *in silico* metagenomes decreases (for example, below five metagenomes), there is an insufficient amount of information to differentiate between low coverage organisms with similar abundances across multiple samples. At four *in silico* metagenomes, BinSanity grouped organisms with similar coverage profiles together, leading to some bins with high redundancy/contamination. Utilizing the refinement script to differentiate bins with high redundancy (as determined by CheckM) using AP clustering and compositional data increased the ARI score from 0.82 to 0.96. Additionally, if over splitting were to occur, a similar refinement step could be used by grouping contigs from these low completion bins into a single file and re-clustering using the refinement script. This method clusters using both coverage and composition information. When using refinement at two *in silico* metagenomes, BinSanity still had the highest ARI at 0.9 compared to 0.83, 0.6, and 0.75 for CONCOCT, MetaBat, and Maxbin, respectively (GroopM could not be run at two *in silico* metagenomes).

### Species Level: diverse-mixture-2

In diverse-mixture-2 (all organisms <10X coverage), BinSanity loses accuracy ( *e.g.* decreased ARI, precision, and V-measure) below 10 *in silico* metagenomes (Figure 5). This trend is associated with a convergence of coverage values across multiple species. BinSanity solely utilizes coverages, therefore it is expected, that as coverages begin to converge, precision will decrease while recall increases (due to contigs from multiple taxa being clustered into the same bins). Similar to diverse-mixture-1, incorporation of the refinement script (refine-contaminated.py) allowed BinSanity to maintain a high ARI above two *in silico* metagenomes; at two metagenomes BinSanity was outperformed by CONCOCT.

**Figure 5.**
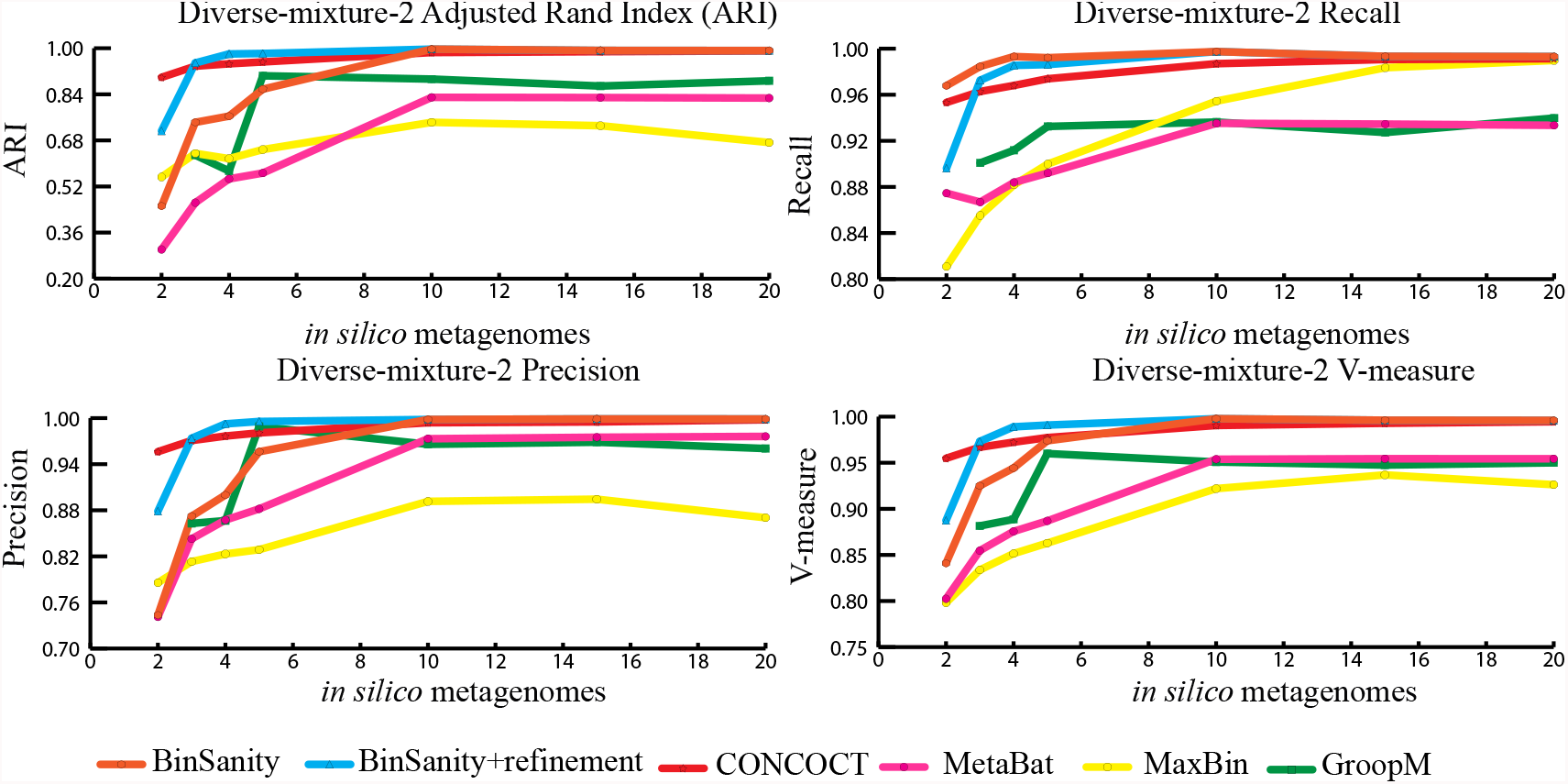
Statistical calculations (bin_evaluation.py) for Adjusted rand index (ARI), Precision, Recall, and V-measure for diverse-mixture-2 which included 50 rare-biosphere organisms. With no refinement BinSanity maintained the highest ARI, Recall, and V-measure above 10 in silico metagenomes (followed closely by CONCOCT). When refinement was incorporated into BinSanity it maintained the highest ARI, Recall, and V-measure above 2 in silico metagenomes. It should be noted that statisticall as recall increases precision decreases and vice versa.

Comparison of CONCOCT, MaxBin, MetaBat, GroopM, BinSanity, and BinSanity+refinement at five *in silico* metagenomes, indicated that BinSanity+refinement produced bins with a higher degree of agreement to the true contig assignments (Figure 6). At five *in silico* metagenomes, (without refinement) BinSanity produced 46 bins compared to the expected 50. When refinement was incorporated into the workflow this number jumped to 54 bins, of which four consisted of small fractions of the target genomes (<6% complete). With refinement, BinSanity was able to accurately split contigs from six organisms that were clustered into two bins. In comparison, CONCOCT, GroopM, MaxBin, and MetaBat produced 70, 92, 40, and 69 bins, respectively. CONCOCT and GroopM produced results with more accuracy, to the expected contig assignment, than MaxBin and MetaBat. GroopM failed to cluster one organism and over split several other organisms. CONCOCT clustered two *Desulfurococcus* species and over split several genomes. MaxBin and MetaBat massively over split genomes and had a high percentage of contigs that were not placed in bins. These results suggest BinSanity can separate low coverage organisms effectively from a large sample set by conducting a first pass using the standard BinSanity script, followed by unsupervised refinement of bins with high contamination and/or low completion. With use of the refinement script, BinSanity maintained a high ARI at three *in silico* metagenome samples, but was surpassed by CONCOCT at two *in silico* metagenomes.

**Figure 6.**
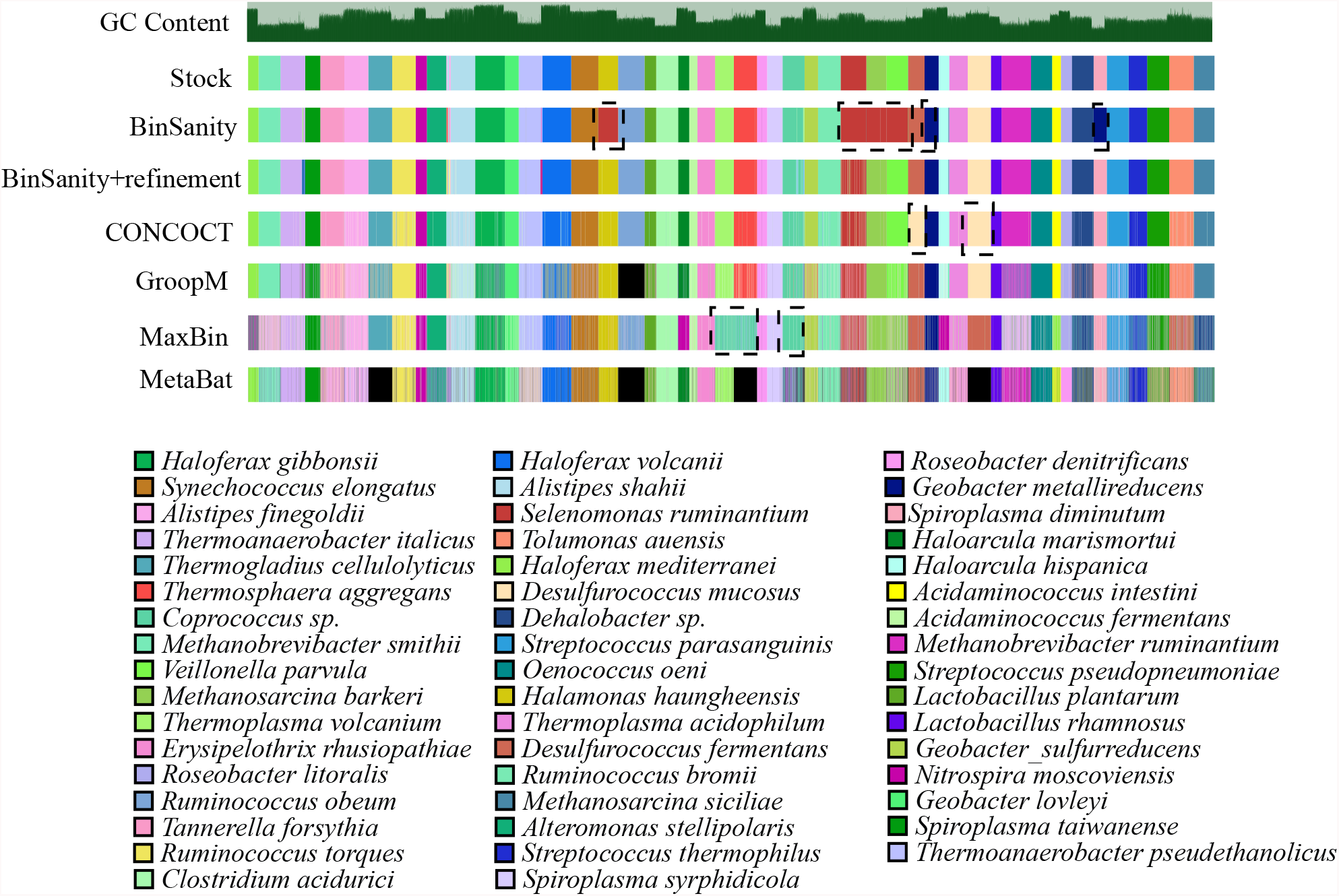
This figure was generated using Anvi’o using a newick tree formating using coverage to order the contigs. This image shows bins output for each of 5 methods, and BinSanity+refinement, at five in silico metagenomes on diverse-mixture-2. Black boxes outline organisms that have been clumped together in different methods. It can be seen that BinSanity without refinement clusters four organisms int one bin 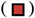 and two organisms into another bin 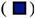. When BinSanity+refinement was used all organisms were able to be teased out. Bins highlighted in CONCOCT are Desulfurococcus fermentans and Desulfurococcus mucosus and were clustered together. The highlighted bins in MaxBin are Thermospaera aggregans and Thermoplasma volcanium. Portions marked in black are contigs that remained unbinned.

### Strain-Level

For the strain-mixture community with 25 organisms (including 4 strains of *Escherichia coli*), BinSanity produced 27 bins using 10 metagenomes. In contrast, CONCOCT, MetaBat, MaxBin, and GroopM produced 32, 41, 18, and 49 bins, respectively, and had lower overall values for the other determined metrics (Figure 7). BinSanity maintained the highest ARI and V-measure regardless of the number of metagenomes used to determine coverage, except at the lowest end of the range (two metagenome). At two *in silico* metagenomes, MetaBat and MaxBin outperformed BinSanity. Most of the methods had difficulty effectively splitting the four closely related *E. coli* strains and two *Escherichia* species (Figure 8 & 9; Supplemental T2-6).

**Figure 7.**
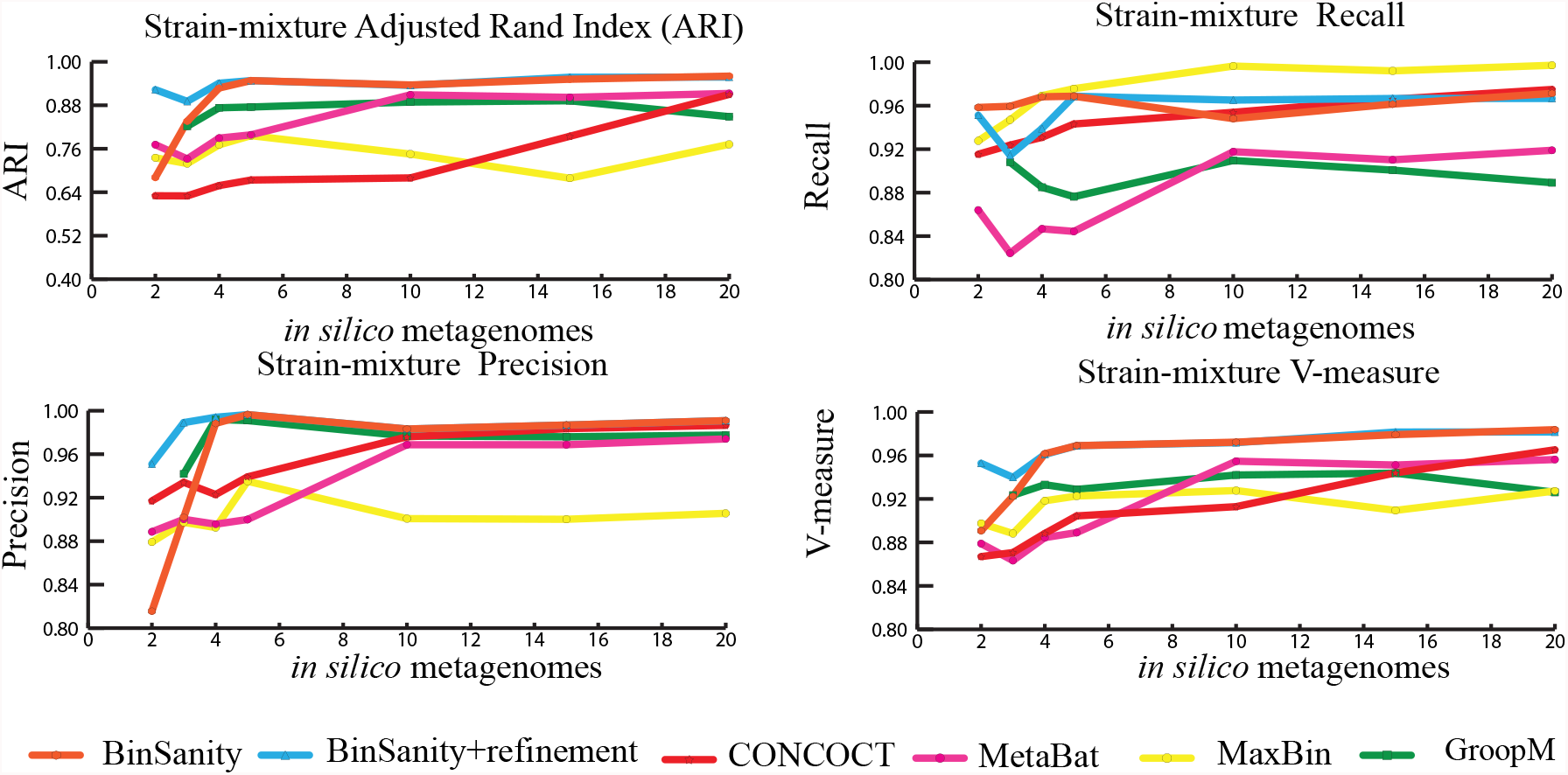
Statistical calculations of Adjusted rand index (ARI), Precision, Recall, and V-measure for the strain-mixture which included 13 abundant biosphere organisms, 12 rare biosphere organisms. Of those 25, four strains of Escherichia coli, and two closlely related Escherichia species are included. BinSanity maintained the highest ARI and V-measure throughout the tests, but similar to other methods saw a drop in accuracy below 4 in silico metagenomes.

**Figure 8.**
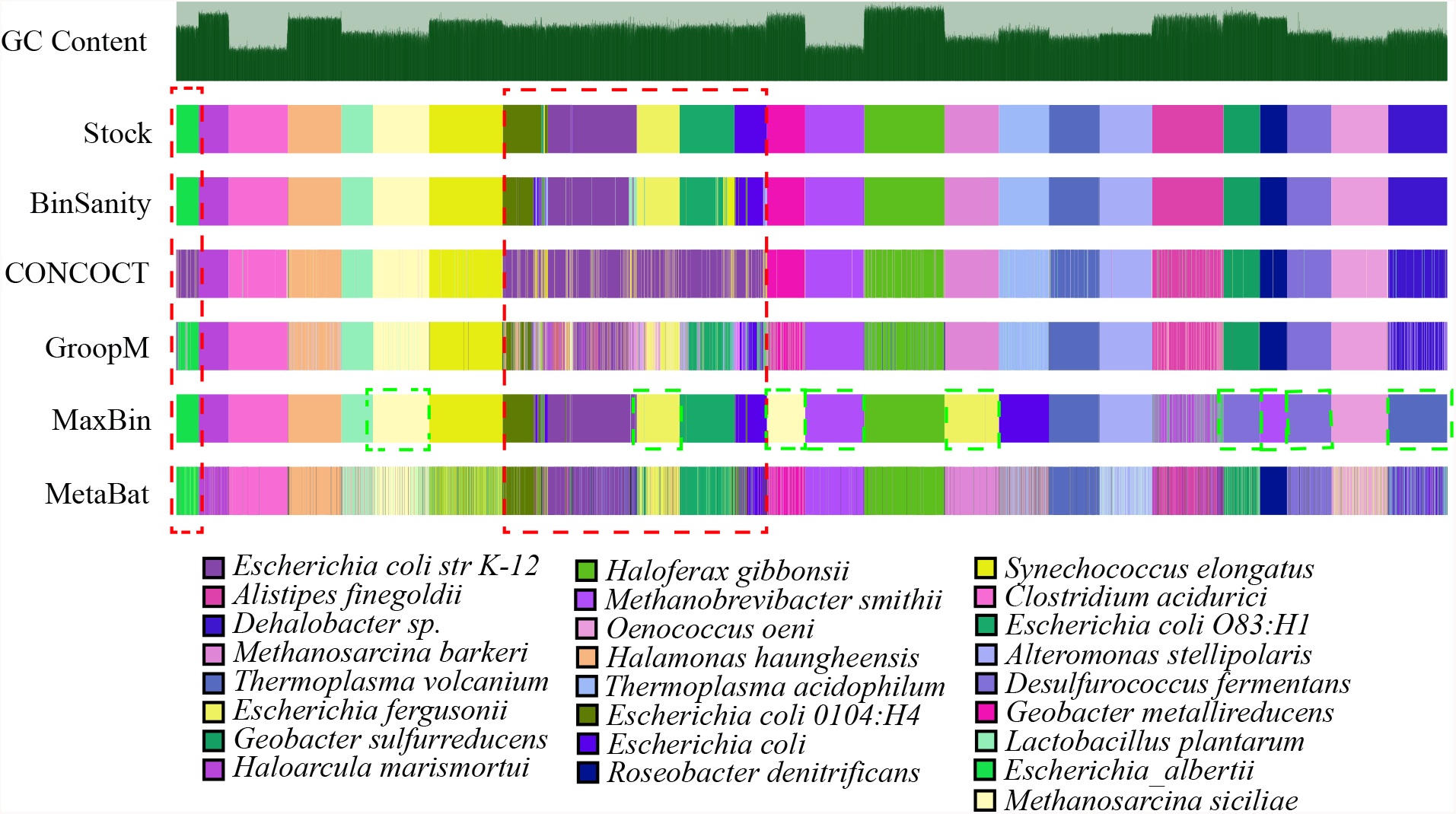
Indicates clustering results for strain-mixture with 25 organisms (4 *Escherichia coli* strains), 2 *Escherichia* genus, and 19 other organisms. The results were visualized using Anvi’o and contigs grouped via coverage and composition. Contigs that were unbinned are emplasized in black. The red box outlines 6 *Escherichia sp.* Green boxes indicate contgis from multiple organisms that have been clumped. Color designations of each organism are based on the stock mixture which represents the actual designations of each contig.

**Figure 9.**
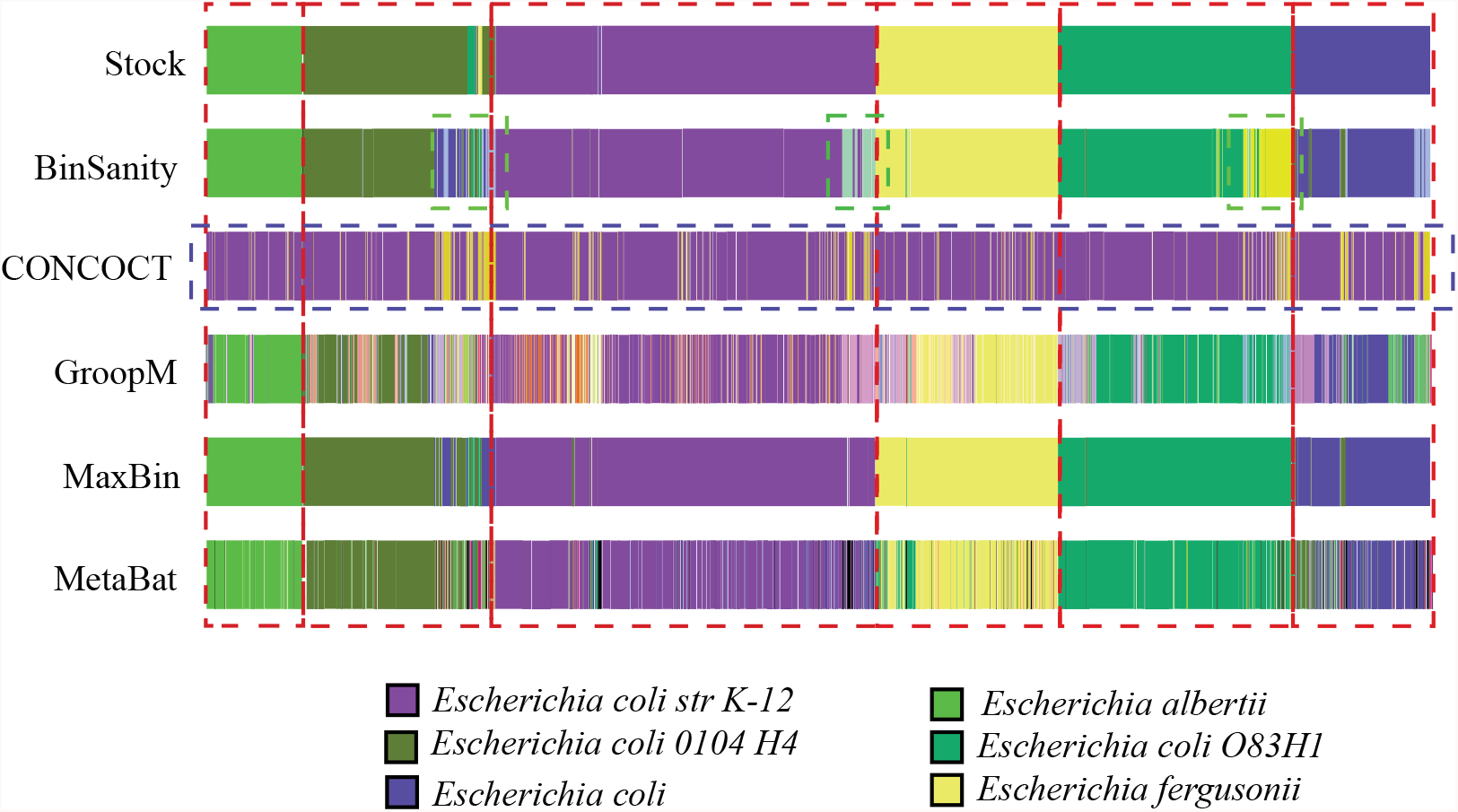
This figure zooms in on the strain-mixture clustering of Escherichia sp. (generated using Anvi’o). The stock row shows the real contig assignments. Black represents contigs that were unclustered. The red boxes outline the bounds of each *Escherichia sp*. The blue box indicates how concoct clustered all of the specie into two bins. The green shows where BinSanity oversplit strains (which corresponds to some of the highly conserved regions of the genome).

At five *in silico* metagenomes, BinSanity, CONCOCT, GroopM, MaxBin, and MetaBat produced 26, 34, 63, 18, and 48 bins, respectively. Of the 26 bins BinSanity produced, 22 were high completion bins with expected redundancy values. The primary difficulty for clustering this dataset for all of the tested methods was accurately differentiating organisms with strain-level similarity. BinSanity generated for four bins from the four *E. coli* genomes. One of the four bins (not included in the 22 high completion genomes mentioned above) had a contamination value of 61.49%, of which 99.5% was due to strain heterogeneity (Bin_18). Bin_18 primarily contained contigs from *E. coli 0104:H4,* but had a large contamination due to the presence of contigs from *E. coli UMN026.* Of the three remaining bins, one contained short contigs <1,500bp in length. The other two were 88% complete bins with between 8% and 13% contamination. One bin contained the remaining contigs from *E. coli* UMN026 that did not incorrectly cluster with *E. coli 0104:H4*, and the other bin contained most of the contigs from *E. coli O83:H1*, with some redundancy due to a small number of contigs containing highly conserved regions from multiple genomes. MaxBin achieved the best resolution of strains, but had difficulty clustering other organisms within the community at a species level (Figure 8 & 9). Metabat and GroopM had a high precision, but an extremely low recall due to high degree of genome splitting. CONCOCT, although approximating the correct results for the other members of the community,largely clustered all 6 *Escherichia* genomes into a single bin.

For the strain-mixture community, GroopM, MetaBat, and MaxBin failed to cluster the most contigs, 261, 56, and 49 contigs, respectively. BinSanity fared better than CONCOCT in accurately representing strains. Based on both the statistics (ARI, precision, and recall) and binning output analysis, BinSanity performed better than the current published unsupervised methods for clustering a community with strain-level variation.

### Infant Gut Metagenome

BinSanity was applied to a metagenomic dataset from a time-series of samples collected from an infant gut environment by Sharon, et al. ^36^ and assembled by Eren, et al. ^37^. The CLC assembled contigs were processed using BinSanity, CONCOCT, GroopM, MaxBin, and MetaBat (Figure 10). The results from the BinSanity method were additionally compared to the output generated by Sharon, et al. ^36^ and Eren, et al. ^37^ (Table 1). The Eren, et al. ^37^ bins were curated using human guided binning via Anvi’o, and Sharon, et al. ^36^ used ESOM^13^ to bin their contigs and manually curated the results. Without use of refinement, BinSanity closely resembled the bins determined manually through Anvi’o^37^. BinSanity split contigs assigned to *Staphylococcus epidermidis* into two bins and clustered contigs assigned to *Propionibacterium acnes* by Anvi’o with *Anaerococcus*. This difference is not observable using the CheckM estimate for completeness, but analysis of the G+C% content indicated that Eren, et al. ^37^ clustered these contigs into two bins, *P. acnes* and *Anaerococcus.* These organisms have divergent compositional makeup, but similar coverage profiles causing BinSanity to group these organisms together. *Candida albicans*, a eukaryote, was difficult to cluster accurately. However, this is to be expected as the task of accurately clustering DNA from eukaryotic genomes is currently beyond the scope of BinSanity and many of the methods discussed in this research.

**Figure 10.**
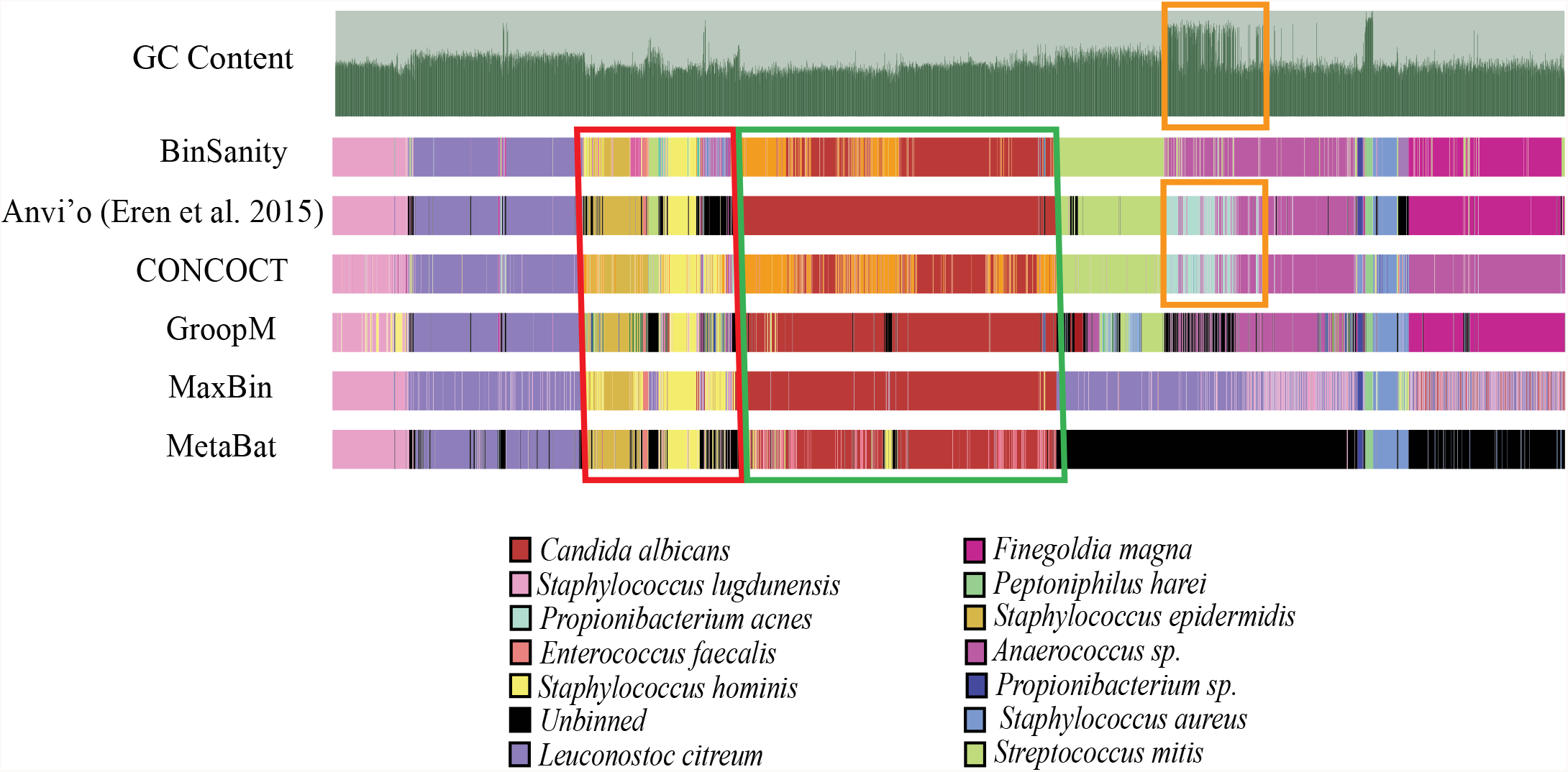
exhibits clustering of 11 co-assembled infant gut metagenomes collected by Sharon et. al (2013) and assembled by Eren et al. (2015) in CLC. Clusters were visualized via Anvi’o. Black indicates contigs that were unbinned. The green box highlight Candida albicans which was clustered into two bins with little loss of completion. Bins highlighted in yellow are the propionibacterium acnes bins found by Anvi’o and CO^-^NCOCT. Bins highlighted in red are staphylococcus strains which each method had difficulty deliminating.

**Table 1.** Shows CheckM calculated compleiton and redundancy percentages for ESOM generated bins curated from the infant gut metagenome collected by Sharon et al. 2013. Percent completion for each organism is compared to BinSanity and Anvi’o curated results. BinSanity and Anvi’o utilized the same inputs of contigs (CLC generated) for clustering while the ESOM generated results used contigs produced via an in house pipeline by Sharon et al. 2013.

| Table 1. Infant Gut Metagenome CheckM comparison<br>(% completion, % redundancy) |  |  |  |
| --- | --- | --- | --- |
| Bin ID | ESOM<br>(Sharon et al.<br>2013) | BinSanity | Anvi'o (Eren<br>et al. 2015) |
| *Staphylococcus aureus* | 99.51 (0.08) | 95.02 (0.66) | 95.02 (0.66) |
| *Staphylococcus lugdunensis* | 84.10 (0.02) | 58.07 (1.72) | 58.07 (1.72) |
| *Staphylococcus epidermidis* | 99.81 (0.00) | 89.06 (0.00) | 90.28 (2.22) |
| *Staphylococcus hominis* | 95.39 (0.57) | 97.26 (2.42) | 97.73 (2.19) |
| *Peptoniphilus harei* | 98.95 (0.00) | 100 (0.00) | 100 (0.00) |
| *Propionibacterium sp.* | 97.86 (0.00) | 98.95 (0.00) | 98.95 (0.00) |
| *Enterococcus faecalis* | 99.25 (0.00) | 99.63 (0.00) | 99.53 (0.00) |
| *Leuconostoc citreum* | 45.64 (0.23) | 62.94 (2.57) | 62.80 (2.57) |
| *Candida albicans* | 34.43 (9.48) | 60.89 (26.92) | 61.76 (27.65) |
| *Finegoldia magna* | 29.25 (0.00) | 32.54 (0.29) | 35.43 (0.60) |
| *Streptococcus mitis* | 16.45 (0.33) | 25.31 (1.00) | 23.10 (0.25) |
| *Propionibacterium acnes* | 5.64 (0.00) | 0 (0.00) | 0.00 (0.00) |
| *Anaerococcus sp.* | 2.51 (0.00) | 11.02 (1.22) | 9.90 (0.00) |
| *Archaea unk* | 0.00 | 6.00(0.00) | 0.00 |

Despite these difficulties, BinSanity closely approximated the manually derived Anvi’o results with higher accuracy than the other unsupervised methods. CONCOCT clustered *Anaerococcus* and *Finegoldia magna*, while creating two highly chimeric bins from four organisms. MetaBat failed to cluster a significant majority of the contigs (69%). MaxBin had difficulty identifying four organisms that were <50% complete and had extremely low contig coverage. GroopM resembled both the BinSanity and Anvi’o results, but overall the bins were less robust and contained less contigs compared to both BinSanity and Anvi’o.

The assembly results from Sharon, et al. ^36^ are not publically accessible (only the contigs assigned to each genome are available) resulting in some variation in the results determined by Sharon, et al. ^36^ and the other methods. These variations can be seen in the *Staphylococcus* bins. For example, *Staphylococcus lugdunensis* was determined to be ∼58% complete by BinSanity, Anvi’o, and CONCOCT (MetaBat 49% complete), but the genome published by Sharon, *et al*. was 84% complete. Overall, BinSanity generated bins reflecting published organisms from this metagenome sampling.

In regards to finding the correct preference value for binning via BinSanity, due to the high variance in coverages in this sample, the preference value was decreased to -10 to account for contigs ranging from 400X to 1X coverage. This induced a higher convergence rate for contigs, increasing the grouping of contigs with similar coverage values. Ultimately, this lead to more complete bins, while larger preference values generated more precise bins but overall had lower recall values.

### A Note on Assigning a Preference Value

Based on results using *in silico* and environmental metagenomic datasets, BinSanity provided a more effective alternative to published unsupervised clustering algorithms. Although BinSanity is an unsupervised method, it is sensitive to changes in the preference value. The preference value sets limits as to how relaxed or stringent AP should be in deciding the number of cluster centers; with low values creating less cluster centers, and high values creating more cluster centers. This preference value is best set by first analyzing the samples in question. For the infant gut metagenome, where a small proportion of contigs had >100X coverage and the remainder had <10X coverage, a preference of -10 was found to be optimal. An initial pass was done at a preference of -5, the BinSanity default value. At this higher preference, BinSanity over split the *Staphylococcus* strains. A similar trend was seen when testing the strain-mixture; a preference of -3000 was used to prevent the over splitting of strains. BinSanity had a tendency to over split bins when strain level variation was detected. Results suggest that this is due to the incorrect mapping of reads sourced from one strain mapping to a related strain and causes coverage values for portions of the genome to deviate from the true mean of the organism. A modification for preference was required when clustering diverse-mixture-2, where coverage values converged and are all <10X. To accurately cluster these samples, a preference of -1 was used induce more splitting.

To predict the correct preference value, it is necessary to look at the range of coverages in a metagenome. When a high range of coverages exists, the preference should be reduced or splitting may occur. When a low range of coverages exists the preference should be increased to prevent mis-clustering of contigs. If a lot of strain-level diversity is expected, preference should be inversely scaled to the number of metagenome replicates ( *e.g.* the more metagenomic samples the lower the preference). Iteratively testing preferences is the best way to find the optimal clustering result while using BinSanity. The affinity-propagation authors state that a good starting point for the preferences is the median or minimum similarity between the most extreme values^16^. Determining an initial preference could be made using the minimum or median similarity of the coverage values seen in a metagenome. When using BinSanity, a tendency towards producing bins with a higher recall can be favored ( *e.g.* using a lower preference) because the refinement function can be used to effectively split high contamination bins. And lastly, BinSanity does not require a set input of all assembled contigs to be effective; as assembly output size increases AP clustering can become computationally prohibitive. Additional methods of grouping contigs prior to BinSanity can be used to simplify this computational step.

## Concluding Remarks

Experimental testing on both real and artificial communities demonstrated that BinSanity outperformed the binning methods CONCOCT, MetaBat, MaxBin, and GroopM when the coverage values for five or more metagenomic samples are available. Additionally, below four metagenomes, when composition information is incorporated via the refinement-function, organisms with similar coverage profiles can be teased apart into accurate genomes. With this refinement step, BinSanity is able to maintain higher precision and recall values compared to the other methods. Based on the unsupervised binning of the infant gut and strain-level communities, BinSanity consistently produced results with higher precision, completeness, and ARI compared to other unsupervised methods. Manually curated results generated similar outcomes, though the time spent manually refining bins can become a limiting factor as the number of MAGs increases. Although preference selection highly impacts the final clustering results for BinSanity, an optimal value can be estimated by factoring in the range of coverage values from the source contigs. Despite the need to manually optimize a preference on an experiment by experiment basis, BinSanity had a higher success at consistently generating accurate genomes from strain- and species-level diversity. The consistency with which BinSanity generates high quality genomes across varying community structures indicates that it is a strong alternative to the compositional based clustering of metagenomic data.

## Software Availability

All scripts used in the generation of these results are available at https://github.com/edgraham/BinSanity/

